# *Streptococcus pneumoniae* serotype 33G: genetic, serological, and structural analysis of a new capsule type

**DOI:** 10.1101/2023.09.11.556596

**Authors:** Sam Manna, Joel P. Werren, Belinda D. Ortika, Barbara Bellich, Casey L. Pell, Elissavet Nikolaou, Ilche Gjuroski, Stephanie Lo, Jason Hinds, Odgerel Tundev, Eileen M. Dunne, Bradford D. Gessner, Stephen D. Bentley, Fiona M. Russell, E. Kim Mulholland, Tuya Mungun, Claire von Mollendorf, Paul V. Licciardi, Paola Cescutti, Neil Ravenscroft, Markus Hilty, Catherine Satzke

## Abstract

*Streptococcus pneumoniae* (the pneumococcus) is a human pathogen responsible for a spectrum of diseases such as pneumonia, sepsis, and meningitis. The capsule is the major pneumococcal virulence factor and is encoded by the capsular polysaccharide (*cps*) locus, a recombination hot spot that has resulted in over 100 distinct capsular polysaccharide types (serotypes) identified to date. Recently, 33X (also known as 10X) was proposed as a putative novel serotype, but the capsule structure had not been elucidated. Here, we provide an in-depth investigation of 33X, demonstrating it is a new pneumococcal capsular serotype. In this study, we screened 12,850 nasopharyngeal swabs from both healthy children and pneumonia patients (adults and children) in Mongolia collected between 2015-2022. We identified 20 pneumococcal 33X isolates. Using whole genome sequencing, we found that the 33X *cps* locus is a chimera of genes from pneumococcal serogroups 35, 10 and 33, as well as other Streptococcal species. Serotyping of 33X pneumococci by the Quellung reaction revealed a unique serological profile, typing as both 10B and 33B. Competitive ELISAs confirmed that antibodies that were generated in mice directed against 33X were inhibited by 33X pneumococci but not 10B or 33B. Lastly, elucidation of the 33X capsule structure revealed that the polysaccharide is distinct from other serotypes, consisting of an O-acetylated hexasaccharide repeat unit of →5)-β-Gal*f*-(1→3)-β-Glc*p*-(1→5)-β-Gal*f* 2Ac-(1→3)-β-Gal*p*NAc-(1→3)-α-Gal*p*-(1→4)-Rib-ol-(5→P→. Therefore, 33X meets the requisite genetic, serological, and biochemical criteria to be designated as a new serotype, which we have named 33G.

**IMPORTANCE:** *Streptococcus pneumoniae* (the pneumococcus) is a bacterial pathogen with the greatest burden of disease in Asia and Africa. The pneumococcal capsular polysaccharide has biological relevance as a major virulence factor, as well as public health importance as it is the target for currently licensed vaccines. These vaccines have limited valency, covering up to 23 of the >100 known capsular types (serotypes) with higher valency vaccines in development. Here, we have characterized a new pneumococcal serotype, which we have named 33G. We detected serotype 33G in nasopharyngeal swabs (n=20) from children and adults hospitalized with pneumonia, as well as healthy children in Mongolia. We show that the genetic, serological, and biochemical properties of 33G differs from existing serotypes, satisfying the criteria to be designated as a new serotype. Future studies should focus on the geographical distribution of 33G and any changes in prevalence following vaccine introduction.

## INTRODUCTION

*Streptococcus pneumoniae* (the pneumococcus) is a common member of the upper respiratory tract microbiota of children. The pneumococcus is also an important human pathogen responsible for a range of diseases such as otitis media, pneumonia, sepsis and meningitis (1). The polysaccharide capsule is a major pneumococcal virulence determinant. The capsule promotes survival within the host by facilitating the evasion of phagocytosis, complement and mucus entrapment (2). Pneumococci exhibit a high level of diversity in capsular polysaccharide structure, with over 100 capsular types (referred to as ‘serotypes’) identified to date (3). Serotyping is the basis for classification of pneumococci. The pneumococcal capsule also has important public health implications as it is the antigenic target for licensed vaccines. Accurate serotyping data are essential in evaluating vaccine impact in carriage and disease, as well as for disease surveillance programs more broadly.

To date, the genomes of over 50,000 pneumococci have been sequenced (4, 5). Sequence analysis has led to the identification of genetic variants of existing serotypes (6–10) as well as new serotypes (3, 11–15). Recently, there has been identification of serotype variants (6, 16) and new serotypes (15) within serogroup 33. The putative new serotype 33X (also known as 10X (17)), relevant to this study, was identified in Thailand (n=5 isolates) (18) and South Africa (n=2 isolates) (16). Although this 33X has been reported previously, little is known about the 33X capsule including whether it is indeed a new serotype or a molecular variant of an existing serotype as the capsule structure has not been investigated. In this study, we identify 33X pneumococci in Mongolia and show the 33X capsule possesses genetic, serological, and structural properties that differ to known serotypes. We name this new serotype 33G.

## RESULTS

### Isolation of 33X pneumococci from Mongolia

In Mongolia, the 13-valent Pneumococcal Conjugate Vaccine (PCV13) was introduced into the paediatric immunization program in a 2+1 schedule (2, 4 and 9 months of age) in a phased manner across districts of the capital city, Ulaanbaatar, from 2016-2018. In 2016 and 2017, three of the districts (Songinokhairkhan, Sukhbaatar and Bayanzurkh) had a catch-up campaign for children aged 3 to 23 months (two doses given two months apart), followed by nation-wide introduction with high coverage (19, 20). Pneumococcal surveillance included the collection of nasopharyngeal swabs from healthy children (cross-sectional surveys in 2015, 2017 and 2022) (21) as well as children (2015–2021) and adults (2019–2022) hospitalized with pneumonia (20, 22, 23) as part of studies assessing the direct and indirect impact of PCV13. Nasopharyngeal swabs (n=12,850) were screened for pneumococci using quantitative PCR (qPCR) targeting the *lytA* gene (24). DNA was extracted from culture-positive samples for molecular serotyping by DNA microarray (25). Overall, 4,800 (37.6%) of the swabs contained pneumococci, and 20 (0.4%) of these contained the putative novel serotype 33X (designated as ‘35A/10B-like’ by DNA microarray). This serotype was detected from 2018 onwards, two years after the phased introduction of PCV13 commenced (2018=3, 2019=6, 2020=7, 2021=0 and 2022=4 isolates) (Supplementary Table S1). The 20 swabs containing 33X pneumococci were from both healthy carriers and patients hospitalized with pneumonia, with no obvious patterns of age, district, or vaccination status (Table 1).

**Table 1.** Participants from which 33X pneumococci were isolated.

| Participant status | Participant age | No. PCV13 doses<sup>a</sup> | District | Year | Isolate name |
| --- | --- | --- | --- | --- | --- |
| Pneumonia | 2 months | 1 | Songinokhairkhan | 2018 | PMP1486 |
| Pneumonia | 9 months | 2 | Songinokhairkhan | 2018 | PMP1488 |
| Pneumonia | 14 months | 3 | Songinokhairkhan | 2018 | PMP1489 |
| Pneumonia | 4 months | 2 | Chingeltei | 2019 | PMP1521 |
| Pneumonia | 2 months | 0 | Songinokhairkhan | 2019 | PMP1596 |
| Pneumonia | 21 months | 2 | Bayanzurkh | 2019 | PMP1597 |
| Pneumonia | 7 months | 2 | Bayanzurkh | 2019 | PMP1598 |
| Pneumonia | 28 years | 0 | Sukhbaatar | 2019 | PMP1609 |
| Pneumonia | 68 years | 0 | Sukhbaatar | 2019 | PMP1614 |
| Pneumonia | 11 months | 3 | Sukhbaatar | 2020 | PMP1592 |
| Pneumonia | 17 months | 3 | Bayanzurkh | 2020 | PMP1593 |
| Pneumonia | 34 months | 2 | Chingeltei | 2020 | PMP1594 |
| Pneumonia | 4 months | 1 | Chingeltei | 2020 | PMP1595 |
| Pneumonia | 34 years | 0 | Chingeltei | 2020 | PMP1612 |
| Pneumonia | 63 years | 0 | Songinokhairkhan | 2020 | PMP1613 |
| Pneumonia | 31 years | 0 | Bayanzurkh | 2020 | PMP1616 |
| Healthy | 5 weeks | 0 | Sukhbaatar | 2022 | PMP1619 |
| Healthy | 17 months | 3 | Songinokhairkhan | 2022 | PMP1620 |
| Healthy | 15 months | 3 | Sukhbaatar | 2022 | PMP1618 |
| Healthy | 16 months | 3 | Songinokhairkhan | 2022 | PMP1617 |

<sup>a</sup> Adults were not eligible for PCV13 vaccination.

### Genetic analysis of the 33X *cps* locus

Using these isolates from Mongolia, we investigated 33X in more detail. A pneumococcal 33X isolate was purified from each swab (Table 1) and whole genome sequencing conducted. We determined genetic lineage using pubMLST (26, 27) (for Multi-Locus Sequence Type) and popPUNK (28) (for Global Pneumococcal Sequencing Cluster). The 33X pneumococci belonged to two genetic lineages ST2754 (GPSC230, n=3) and ST6318 (GPSC687, n=17).

The *cps* locus of the 33X isolates from Mongolia has over 99.7% DNA sequence identity to the 33X *cps* sequences first reported by van Tonder et al (18) as well as by Mostowy et al (17) where it was referred to as 10X. These *cps* sequences were previously identified in isolates from Thailand (ST5123, GPSC293) (18) and South Africa (ST5178, GPSC241) (16). We examined the 33X *cps* locus of the 20 isolates from Mongolia using DNA sequence alignment and genome annotation by RAST (29), finding they have the same mosaic structure as the previously described 33X *cps* locus, comprised of genes with high DNA identity to sequences from serogroups 35 (*wzg, wzh, wzd, wze, wchA, wciB)* (Figure 1A), 10 (partial *wzy* sequences*, wcrB, wcrC, wcrD, wciF*) and 33 (*wzx*, *wciG* and *glf*) (Figure 1B).

**Figure 1.**
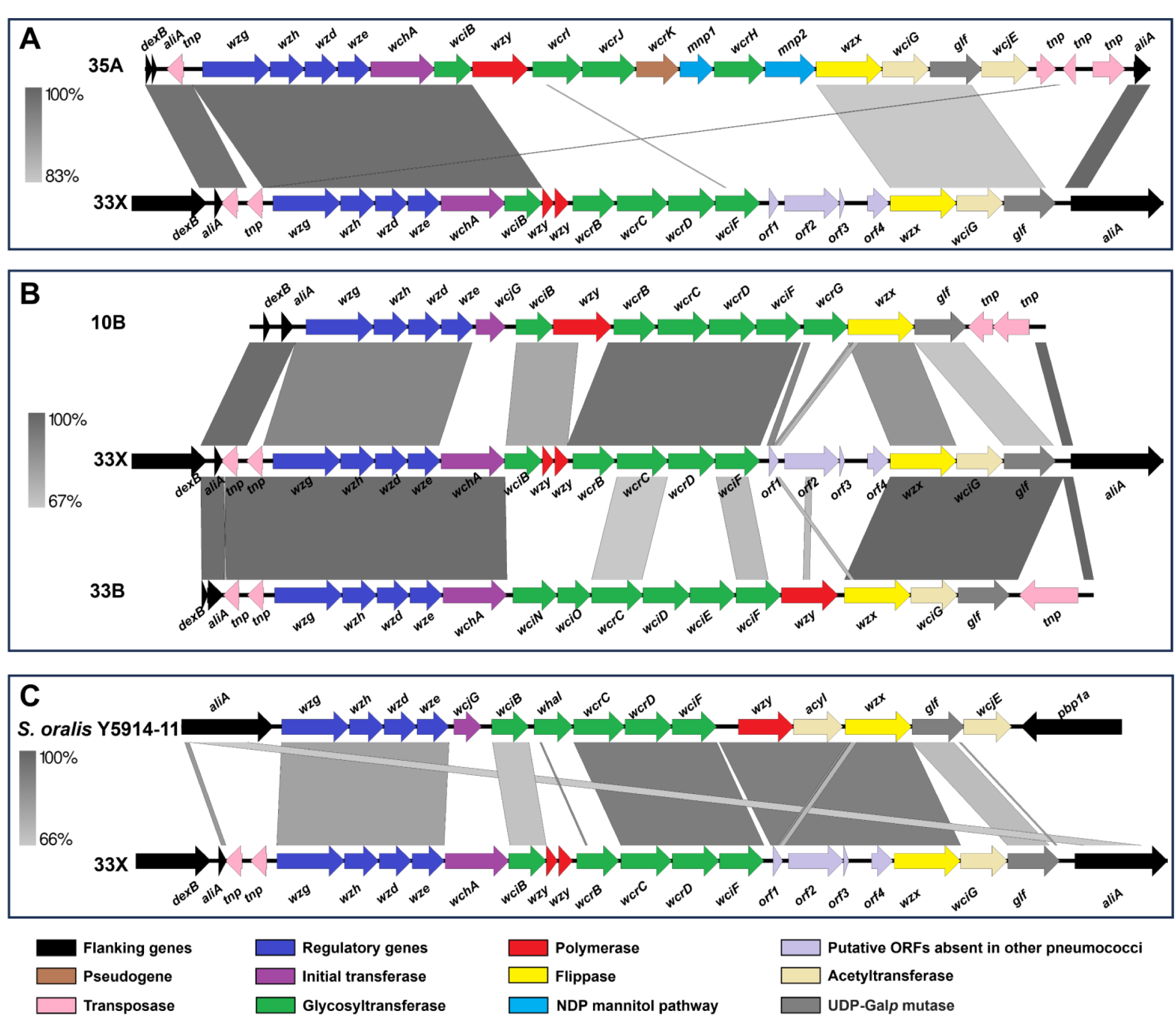
Schematic representation of the 33X *cps* locus with the 35A (A), 10B and 33B (B) *cps* reference sequences (48) and *S. oralis* subsp. *dentisani* strain Y5914-11 *cps* locus (Genbank accession no. NZ_NCUW01000028) (30). Images were generated using Easyfig version 2.2.5 (73).

The 33X *cps* locus contains four putative open reading frames. *Orf1* exhibits partial similarity to part of the *wcrG* and *wzx* genes from the serotype 10B reference *cps* locus (Genbank accession CR931650), possibly representing 10B remnant sequences following a recombination event at this site. Interestingly, *orf2, orf3* and *orf4* do not show sequence similarity to any pneumococcal *cps* genes. Rather, they show similarity to a region in the *cps* locus from *Streptococcus oralis* subsp. *dentisani* strain Y5914-11 (Figure 1C), a blood isolate from a patient with infective endocarditis from Denmark (30). Between *wciB* and *wcrB* are two sequences with similarity to part of the *wzy* gene, which encodes the polymerase from the serotype 10B reference *cps* locus. However, the deletions apparent in these sequences suggest these would not encode a functional enzyme. *Orf2* is predicted to encode a Wzy homolog, with the predicted amino acid sequence exhibiting high identity to O-antigen ligase family proteins from *S. oralis* (97.2%, Genbank accession WP_204956359) and *Streptococcus intermedius* (97.6%, Genbank accession MBF1713530). Among pneumococcal Wzy proteins, ORF2 exhibits the highest identity to Wzy from serotype 37 (41.7%, Genbank accession CAI34388). Lastly, ORF3 and ORF4 are partial sequences that match portions of the *S. oralis* acyltransferase (86.5% and 74.7%, to *S. oralis* acyltransferase Genbank accession WP_084971822 for ORF3 and ORF4, respectively). However, the *orf3-orf4* region in the 33X *cps* locus contains multiple frameshift mutations suggesting it is unlikely to encode a functional acyltransferase (Supplementary Figure S1).

To determine the implications of this genetic variation for molecular approaches for serotyping, we tested the 33X isolates in five different whole genome sequencing-based serotyping tools (PneumoCaT (31), seroBA (32), SeroCall (33), PneumoKITy (34) and PfaSTer (35)) as well as DNA microarray (as described above) and the TaqMan Array Card, a qPCR-based assay (36). Most tools were unable to designate a serotype to 33X pneumococci (Supplementary Table S2). Tools that report the closest ‘hit’ (PneumoCaT, PneumoKITy and PfaSTer), were matches to serotypes in serogroup 35. As described earlier, DNA microarray designates 33X as 35A/10B-like due to the detection of *cps* genes from serotypes 35A and 10B. The TaqMan Array Card mistyped these isolates as 10B. The seroBA tool was the only one to detect 33X (designated as 10X in this tool) given the *cps* locus from the isolates previously identified in Thailand and South Africa is included in the seroBA database.

### Serological properties of 33X pneumococci

To investigate the serological properties of the 33X capsule, the isolates from Mongolia were examined using the Quellung reaction, the gold standard for pneumococcal serotyping (37, 38) using commercially available typing sera from the Statens Serum Institut (SSI). All isolates reacted with both group 10 and 33, but not group 35, sera. When tested against the relevant factor sera, the 33X pneumococci reacted with 10b, 10d and 33f factor sera. Thus, the 33X pneumococcal isolates serotyped as both 10B and 33B by Quellung, despite being confirmed as pure cultures by whole genome sequencing and the 10B and 33B control strains reacting as expected (Table 2).

**Table 2.** Quellung reaction profile of 33X pneumococci with ‘+’ and ‘-’ denoting a positive and negative reaction with the typing sera, respectively.

|  | Sera | 33X | 10B | 33B |
| --- | --- | --- | --- | --- |
| **First pool** | **A** | - | - | - |
|  | **B** | - | - | - |
|  | **C** | - | - | - |
|  | **D** | - | - | - |
|  | **E** | + | + | + |
|  | **F** | - | - | - |
|  | **G** | - | - | - |
|  | **H** | - | - | - |
|  | **I** | - | - | - |
| **Second pool** | **P** | - | - | - |
|  | **Q** | - | - | - |
|  | **R** | - | - | - |
|  | **S** | + | + | - |
|  | **T** | **+** | **-** | **+** |
| **Group** | **10** | **+** | **+** | **-** |
|  | **12** | **-** | **-** | **-** |
|  | **21** | **-** | **-** | **-** |
|  | **35** | **-** | **-** | **-** |
|  | **33** | **+** | **-** | **+** |
|  | **39** | **-** | **-** | **-** |
| **Factors – Group 10** | **10b** | **+** | **+** | **-** |
|  | **10d** | **+ (weak)<sup>a</sup>** | **+** | **-** |
|  | **10f** | **-** | **-** | **-** |
| **Factors – Group 33** | **33b** | **-** | **-** | **-** |
|  | **33e** | **-** | **-** | **-** |
|  | **33f** | **+** | **-** | **+** |
|  | **6a** | **-** | **-** | **-** |
|  | **20b** | **-** | **-** | **-** |

<sup>a</sup> Reaction is weak with one 1 µl loopful of 10d factor sera. A stronger reaction was obtained using two 1 µl loopfuls

To investigate the immunological properties of 33X isolates, we conducted competition ELISAs. To obtain sera, we infected five-day old C57BL/6 mice with a 33X pneumococcal strain (PMP1486) to establish nasopharyngeal colonization and collected serum seven weeks post-infection. The serum was pre-incubated with pneumococci expressing serotype 33X, 10B, 33B, or 14 capsules as competitors before conducting the 33X ELISA. Pre-incubation of serum with 33X pneumococci resulted in greater inhibition (76.6%) compared with pre-incubating with 10B (36.4%), 33B (33.9%) or 14 (negative control, 28%), suggesting that the immunological properties of 33X differ to 10B and 33B (Figure 2).

**Figure 2.**
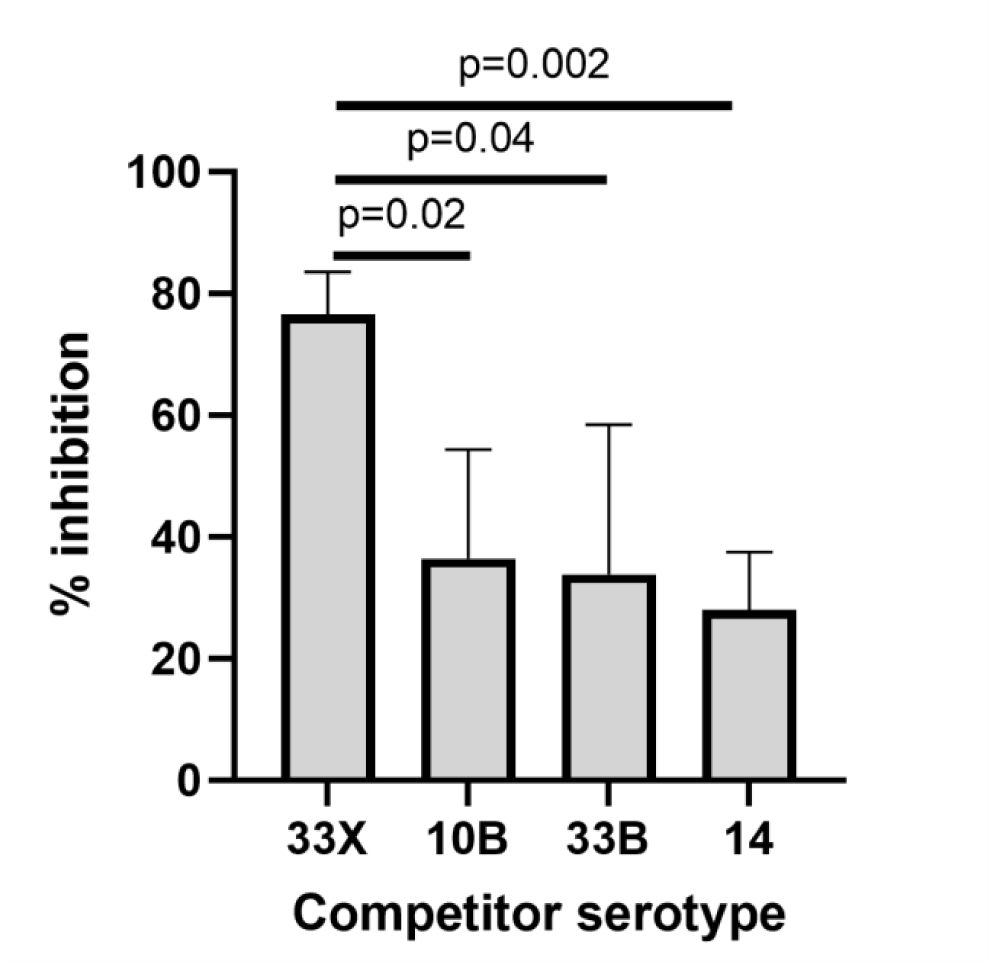
Competition ELISA with serum from mice colonized with 33X pneumococci pre-incubated with serotypes 33X, 10B, 33B or 14 as a competing capsular antigen source. Data presented are mean + standard deviation and analysed by unpaired t test.

Taken together, these data show that 33X pneumococci have genetic and immunological properties that are distinct from previously characterised pneumococcal serotypes.

### Elucidation of 33X capsule repeat unit structure

We next determined the repeat unit structure of the polysaccharide purified from 33X isolates by use of chemical analysis and nuclear magnetic resonance (NMR) spectroscopy. Acid hydrolysis of the serotype 33X polysaccharide followed by derivatisation and gas chromatography-mass spectrometric (GLC-MS) analysis showed the presence of ribitol (Rib-ol), glucose (Glc), galactose (Gal) and N-acetylgalactosamine (GalNAc) in the molar ratio of 1.6:1.0:2.2:0.8. Performing the reduction step with NaBD_4_ and the absence of a deuterium label in the ribitol derivative confirmed that ribitol was not derived from ribose. Composition analysis following methanolysis gave comparable results and confirmed the absence of uronic acid in the repeat unit. Determination of the glycosidic linkages was achieved by GLC-MS analysis of the partially methylated alditol acetates which identified: 4,5-Rib-ol, 3-Glc*p*, 4-Gal*p* or 5-Gal*f*, 3-Gal*p*, and 3-GalNAc*p* in the molar ratio of 0.4:1.6:2.1:1.0:0.7. These data are consistent with a linear hexasaccharide repeat unit without branch points and side chains, unlike the repeat units reported for related serotypes 10B (39) and 33B (40).

The ^1^H NMR spectrum of the serotype 33X polysaccharide (Figure 3A) at 25°C (500 MHz) showed five signals and an <u>H</u>-COAc peak at 4.93 ppm in the anomeric region, ring protons and peaks from low molecular weight components (glycerol, buffer, and amino acids).

**Figure 3.**
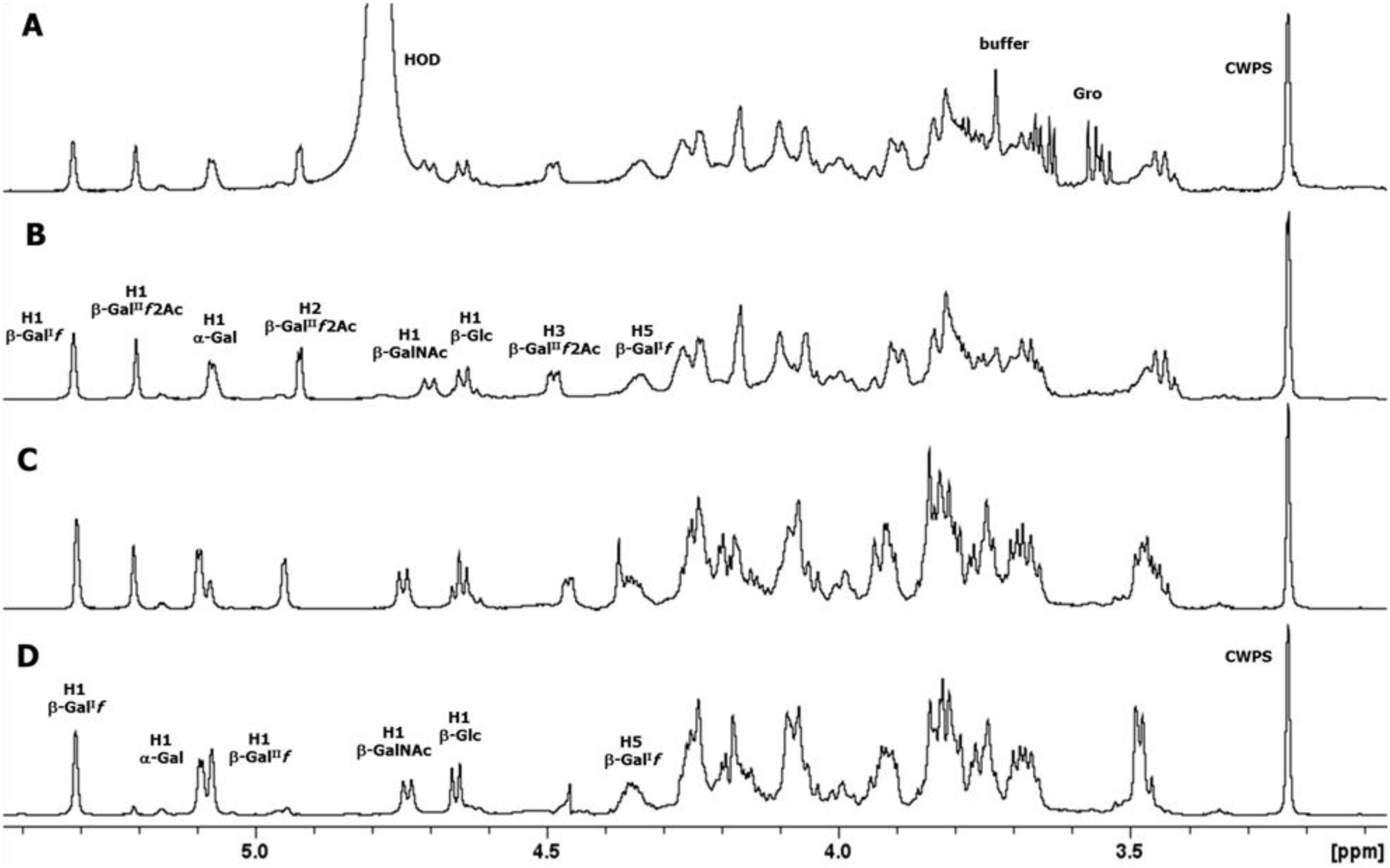
Expansion of the 1D proton NMR spectra of pneumococcal serotype 33X polysaccharide, some diagnostic anomeric and ring protons are labeled. (A) 1D ^1^H NMR at 500 MHz and 25°C, (B) 1D DOSY at 500 MHz and 25°C, (C) 1D DOSY at 600 MHz and 70°C (partially de-O-acetylated), and (D) 1D DOSY at 600 MHz and 60°C (de-O-acetylated polysaccharide).

The full spectrum (Supplementary Figure S2A) contains an O-acetyl signal at 2.14 ppm and an N-acetyl signal at 2.03 ppm indicating that serotype 33X polysaccharide has an O-acetylated hexasaccharide repeating unit containing an N-acetyl sugar. The peaks from HOD (water), process residuals and amino acids were readily removed in the diffusion ordered spectroscopy (DOSY) experiment in Figure 3B (and Supplementary Figure S2B) to give the polysaccharide signals together with peaks from residual cell wall polysaccharide (CWPS) with the characteristic choline resonance at 3.23 ppm. Further NMR studies were performed at 600 MHz and at higher temperatures that resulted in gradual de-O-acetylation (Figure 3C) leading to the NMR spectrum of the polysaccharide backbone shown in Figure 3D (and Supplementary Figure S2D). The structure of the 33X hexasaccharide repeat unit was determined on the de-O-acetylated sample by use of an array of ^1^H-^1^H homonuclear experiments (COSY, TOCSY and NOESY) and ^1^H-^13^C heteronuclear correlation experiments (HSQC and HMBC) described in the Supplementary Figures S3-5. The ^1^H-^31^P HMBC experiment (Supplementary Figure S6) showed that the phosphodiester signal at 0.34 ppm was correlated to the H5s of ribitol and the broad proton signal at 4.35 ppm, assigned to H5 of β-Gal^I^*f*. The proton/carbon pairs are labeled in the HSQC spectrum (Figure 4), the NMR data are collected in Table 3.

**Figure 4.**
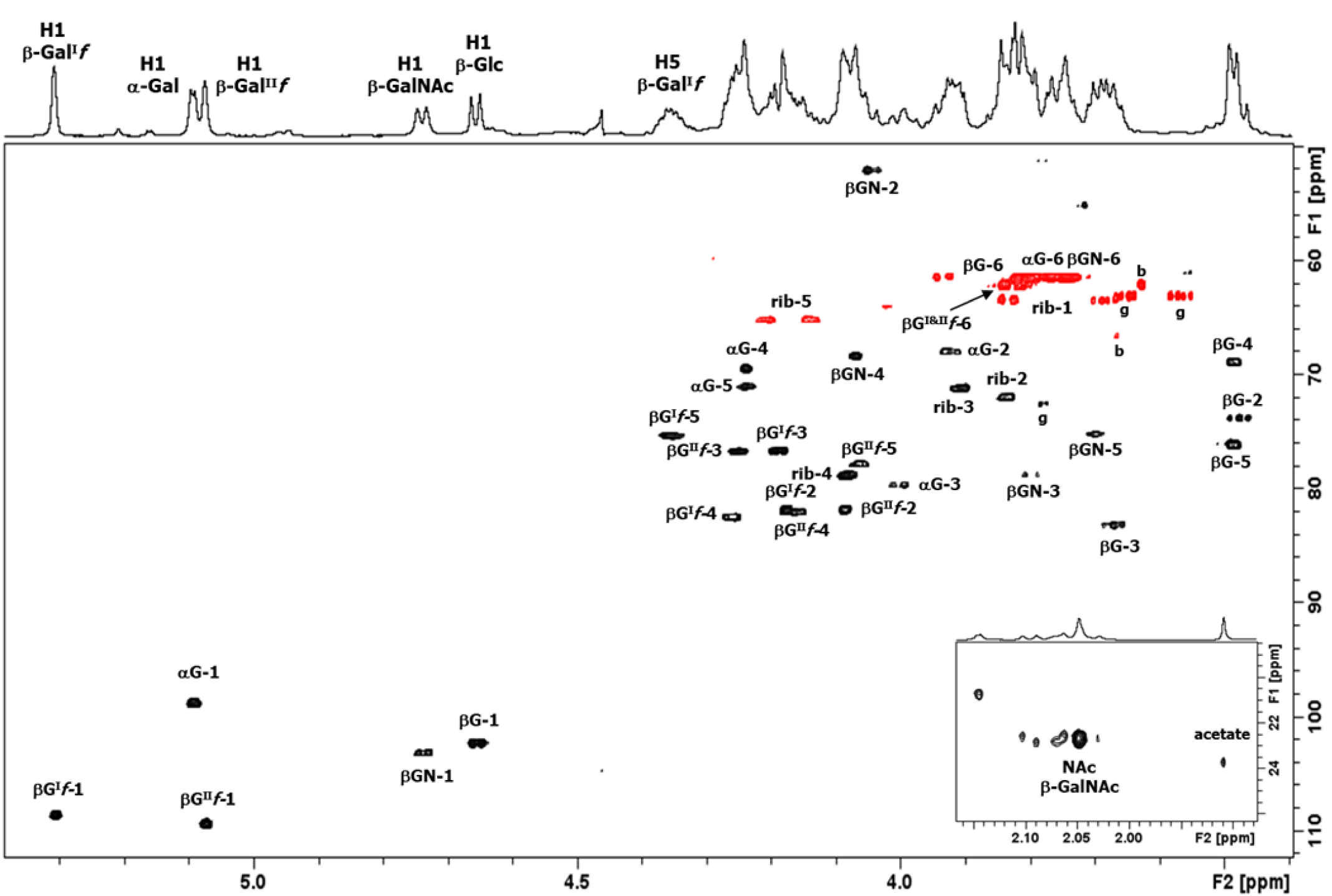
Expansion of the HSQC spectrum of polysaccharide 33X recorded at 600 MHz (60°C), the crosspeaks from the methyl region of the spectrum are shown in the inset. All the hexasaccharide repeat unit (including ribitol-P) proton/carbon crosspeaks have been labeled according to the carbon atom of the corresponding residue (βG^I^*f* = β-Gal^I^*f*, αG = α-Gal*p*, βG^II^*f* = β-Gal^II^*f*, βGN = β-GalNAc, β-G = β-Glc and rib = ribitol). Additional peaks are due to buffer (b) and glycerol (g).

**Table 3.**
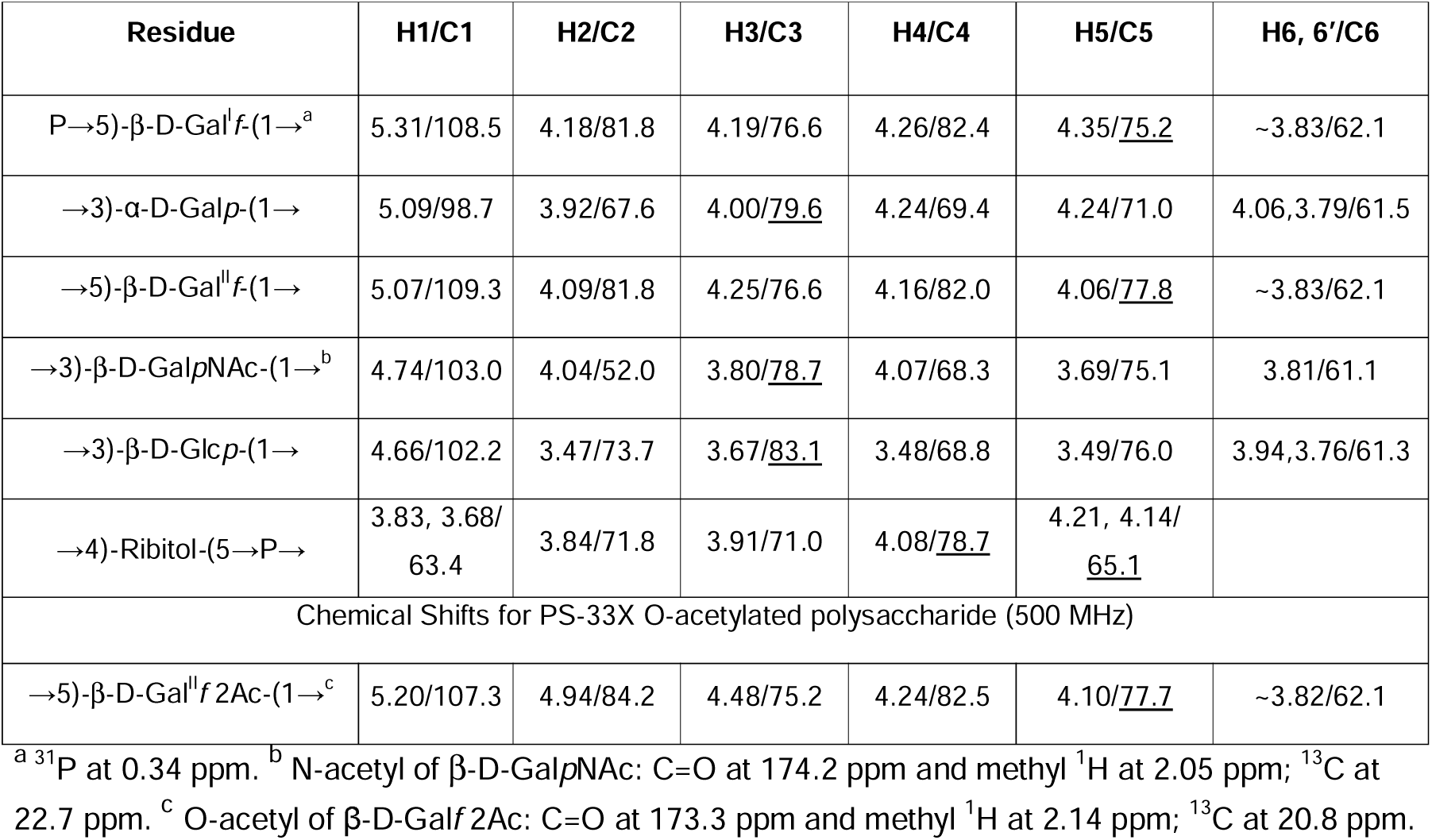
NMR Chemical Shift for serotype 33X polysaccharide (de-O-acetylated) and O-acetylated Gal^II^*f* (lower panel). Linkage carbons are underlined.

Finally, the linkages and sequence of sugar residues followed from the ^1^H-^13^C HMBC inter-residue correlations are labeled in Figure 5. Thus, NMR analysis established the structure of the hexasaccharide repeat unit of serotype 33X as →5)-β-Gal^I^*f*-(1→3)-β-Glc*p*-(1→5)-β-Gal^II^*f*-(1→3)-β-Gal*p*NAc-(1→3)-α-Gal*p*-(1→4)-Rib-ol-(5→P→.

**Figure 5.**
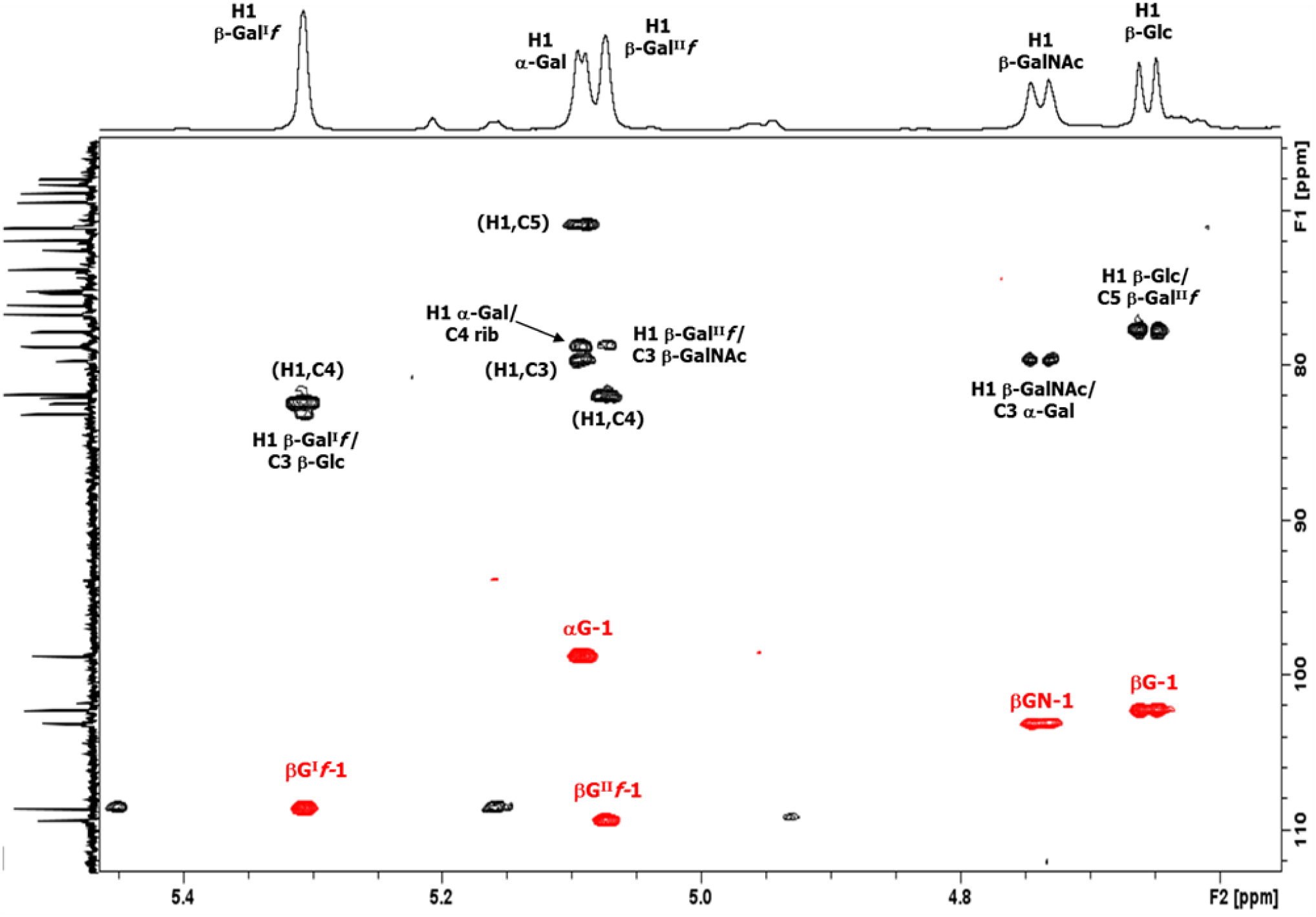
The 2D ^1^H-^13^C overlay of the anomeric region for polysaccharide 33X: HSQC-DEPT (red)/HMBC (black) recorded at 600CMHz (333 K). The intra- and inter-residue HMBC crosspeaks from H1 are labeled according to the carbon atom of the corresponding residue (βG^I^*f* = β-Gal^I^*f*, αG = α-Gal*p*, βG^II^*f* = β-Gal^II^*f*, βGN = β-GalNAc, β-G = β-Glc and rib = ribitol). The HMBC experiment was optimized for a coupling constant of 6CHz and established the following linkages: H1 of β-Gal^I^*f* to C3 of β-Glc, H1 of α-Gal*p* to C4 of ribitol, H1 of β-Gal^II^*f* to C3 of β-GalNAc, H1 of β-GalNAc to C3 of α-Gal*p* and H1 of β-Glc to C5 of β-Gal^II^*f*.

### Identification of O-acetylation at C2 of β-Gal^II^f

NMR experiments performed on the fully and partially O-acetylated polysaccharide (Figures 3A and 3C) confirmed the presence of 5-linked β-Gal^II^*f* 2Ac with diagnostic signals for H1 at 5.20, H2 at 4.93 and H3 at 4.48 ppm, and the O-acetyl singlet at 2.14 ppm. TOCSY of the partially O-acetylated polysaccharide elucidated the proton spin systems of both 5-linked β-Gal^II^*f* 2Ac and the de-O-acetylated 5-linked β-Gal^II^*f* residues (Supplementary Figure S7). Finally, HSQC-DEPT and HMBC of the fully O-acetylated polysaccharide, displayed as an overlay in Figure 6, identified the carbon resonances for this residue. In addition, the inter-residue correlations confirmed the sequence and linkages of the serotype 33X O-acetylated hexasaccharide repeat unit. The NMR assignments for the 5-linked β-Gal^II^*f* 2Ac residue are provided in Table 3, lower panel.

**Figure 6.**
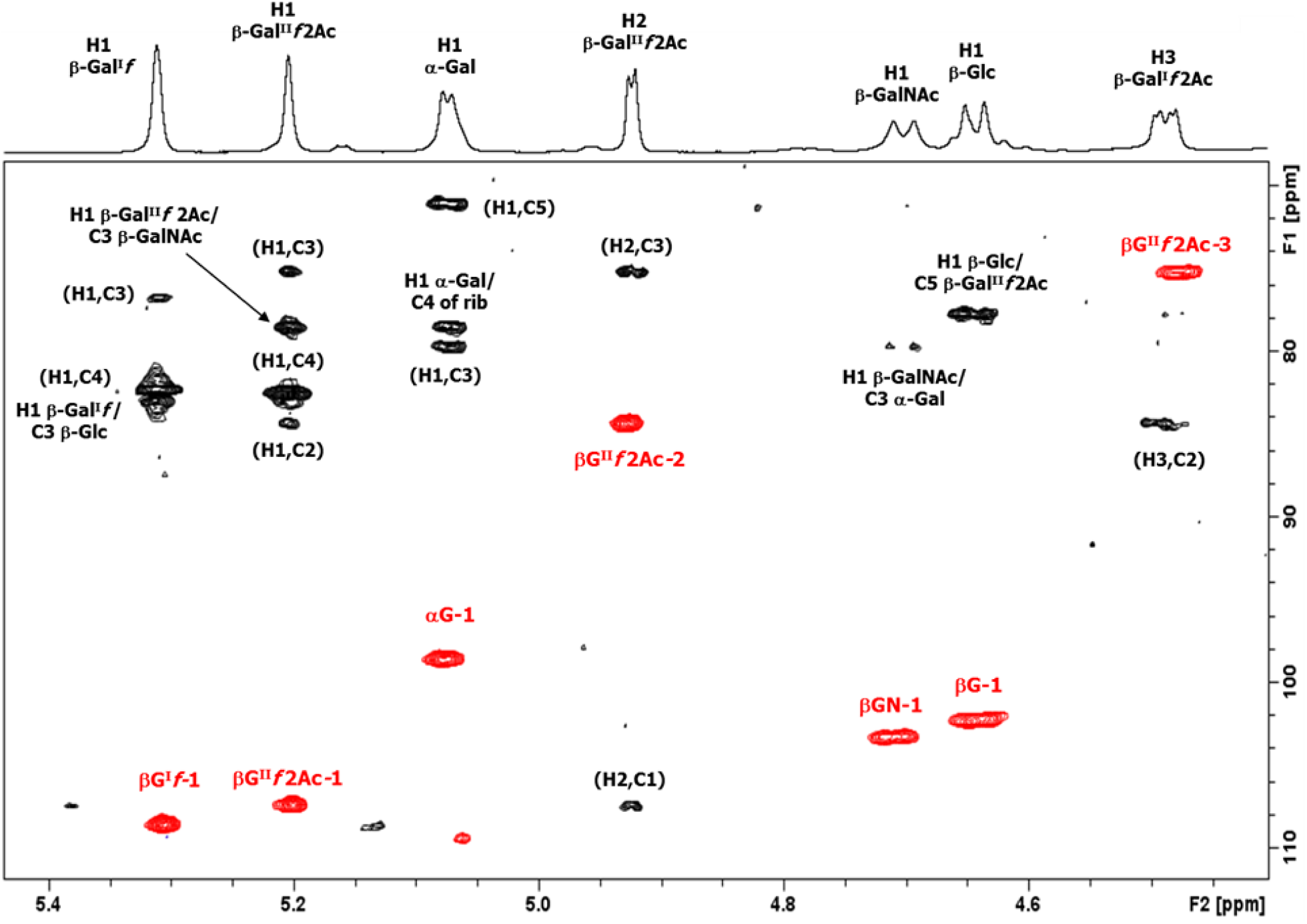
The 2D ^1^H-^13^C overlay of the anomeric region for O-acetylated polysaccharide 33X: HSQC-DEPT (red)/HMBC (black) recorded at 500CMHz (298 K). The intra- and inter-residue HMBC crosspeaks from diagnostic anomeric proton signals are labeled according to the carbon atom of the corresponding residue (βG^I^*f*, = β-Gal^I^*f*, αG = α-Gal*p*, βG^II^*f* 2Ac= β-Gal^II^*f* 2Ac, βGN = β-GalNAc, β-G = β-Glc and rib = ribitol). The HMBC experiment was optimized for a coupling constant of 6 Hz.

Overall, the repeat unit of 33X is an O-acetylated hexasaccharide with components in common with both the 10B and 33B repeat units, but not identical to either. Based on their known enzymatic activities (41), we were able to assign a function to all the enzymes encoded in the 33X *cps* locus to the linkages in the repeat unit, further supporting the proposed structure and its designation as a new pneumococcal serotype, hereby referred to as 33G (Figure 7).

**Figure 7.**
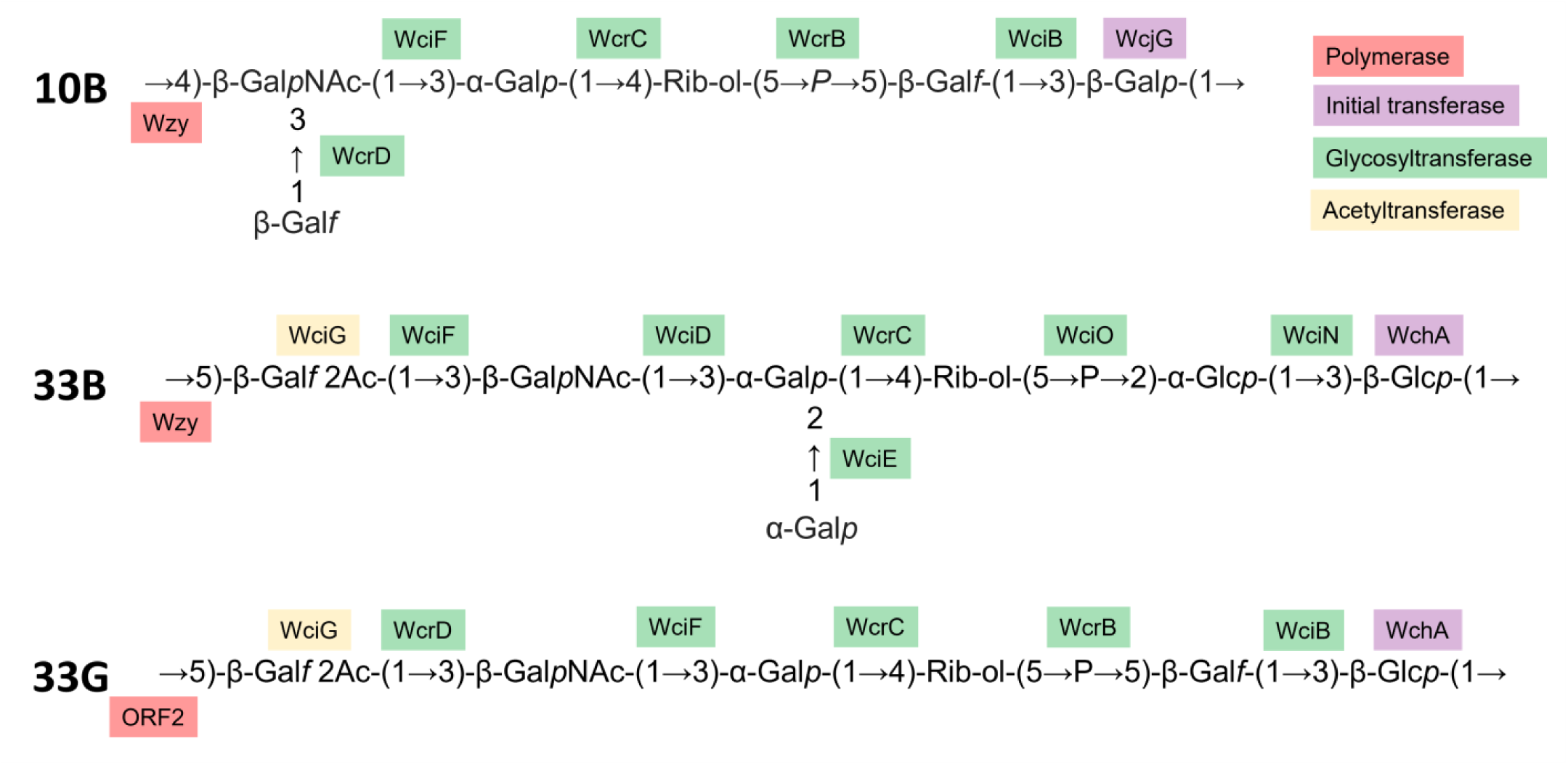
Polysaccharide repeat unit structure of 10B (74), 33B (40), and new serotype 33G (this study), as well as proposed enzymatic assignments based on the literature (15, 41).

## DISCUSSION

Traditionally, pneumococci are classified into serotypes based on differences in serological reactions using the Quellung reaction (42–44). Different serotypes produce a distinct capsular polysaccharide (44–47) encoded by a unique *cps* locus (48). Previously, seven isolates from Thailand (17, 18) and South Africa (16) with the serotype designation 33X (also known as 10X) were identified. No investigation had been conducted into the structural properties of the capsule and it was therefore not known whether 33X is a novel pneumococcal serotype. In this study, we provide a comprehensive investigation of this putative new serotype using 20 isolates that we identified in Mongolia.

Consistent with the previous reports (17, 18), the 33X isolates from Mongolia possess a unique *cps* locus compared with known serotypes. Using the gold standard Quellung reaction, we show that 33X pneumococci exhibit a unique serological profile, typing as both 10B and 33B. Importantly, pre-incubation of serum from mice colonized with 33X pneumococci resulted in greater ELISA inhibition than pre-incubation with either 10B or 33B pneumococci. Serotype 35A was not explored further given the negative Quellung reaction with group 35 sera. Together, these data indicate that serological responses to 33X are distinct from the related serotypes 10B and 33B. Using 1D and 2D NMR and sugar composition analyses, we interrogated the 33X capsule, demonstrating a structure that possessed features of both 10B and 33B, yet was not identical to either nor any other known pneumococcal serotype. Taken together, these data provide the requisite genetic, serological, and biochemical evidence that 33X is a new pneumococcal serotype, hereby referred to as 33G.

We have proposed placing this new serotype in serogroup 33 rather than 10 since five of residues in the 33G hexasaccharide repeat unit are conserved in 33B compared with four for 10B. Like 33B, 33G and other serotypes in serogroup 33 (33C and 33D) also possess a hexasaccharide backbone containing ribitol-5-P. In addition, 33G has an O-acetylated β-Gal*f* residue, found in all members of serogroup 33 and lacking in serogroup 10 capsules (39, 40). The presence of O-acetyl groups in the 33G capsule is likely to have important biological relevance given that O-acetylation impacts the structure of the polysaccharide, and is an epitope in pneumococcal capsules readily recognised by the immune system (44, 49). Such immune pressure has led to the emergence of new serotypes that lack the O-acetyl modification (15, 50–52). Lastly, a characteristic feature found in all serotypes in serogroup 10 is a terminal side chain β-Gal*f* (11, 39), which is lacking in 33G.

The 33G *cps* locus is a mosaic of genes from multiple sources including three pneumococcal serogroups (35, 10 and 33) as well as other streptococci. The emergence of such a complex *cps* is the result of several recombination events. The pneumococcal *cps* locus is a known recombination hotspot and examples of recombination with the *cps* locus of other pneumococci have been reported previously (6, 17). Similar recombination events with non-pneumococcal streptococci also occur, as exemplified by the recently described serotype 10D, which contains capsular genes from serotypes 6C, 39 and oral streptococci (11). The earliest detection of 33G was in an invasive disease isolate from South Africa in 2007 (16). However, with limited epidemiological data and small number of isolates (n=27 in total across three countries including the present study), it is difficult to determine when and where 33G emerged.

The chemical and NMR studies performed on the serotype 33G polysaccharide elucidated the O-acetylated repeat unit as →5)-β-Gal^I^*f*-(1→3)-β-Glc*p*-(1→5)-β-Gal^II^*f* 2Ac-(1→3)-β-Gal*p*NAc-(1→3)-α-Gal*p*-(1→4)-Rib-ol-(5→P→. This structure explains the cross-reactivity observed with serotypes 10B and 33B by Quellung, due to the common structural features seen between these serotypes. Serotypes 10B and 10F contain the tetrasaccharide β-Gal*p*NAc-(1→3)-α-Gal*p*-(1→4)-Rib-ol-(5→P→5)-β-Gal*f*-(1→, but the β-Gal*p*NAc is a branch point 3,4-linked and 4,6-linked respectively, not 3-linked as in 33G. Serotypes 33B and 33D contain the pentasaccharide →3)-β-Glc*p*-(1→5)-β-Gal*f* 2Ac-(1→3)-β-Gal*p*NAc-(1→3)-α-Gal*p*-(1→4)-Rib-ol-(5→P→, but the α-Gal*p* is a branch point with a terminal α-Gal*p* group on C-2 (40), not only 3-linked as in 33G. Additionally, the large deletions in the *wzy* gene in the 33G *cps* locus suggest it is unlikely to encode a functional polymerase. However, *orf2* is predicted to encode a *wzy* homolog, suggesting the acquisition of ORF2 from another streptococcal species has replaced the ancestral Wzy as the polymerase responsible for linking the 33G repeat units together.

Our findings have implications for surveillance using both phenotypic and genotypic serotyping methods. For researchers relying on serological typing methods, like the Quellung reaction, 33G will yield two serotype results (10B and 33B). Therefore, there is potential for 33G to be mistyped as 10B or 33B (or even 10F in some cases if the operator misses the weak reaction with 10d factor sera). Interestingly, 33G pneumococci did not react with any of the sera recognising serogroup 35, despite a large portion of the 33G *cps* locus with high homology to serogroup 35. This likely occurred because the serogroup 35-derived genes include the transcriptional regulatory region (*wzg-wze*), the initial transferase (*wchA*) and one glycosyltransferase (*wciB*), which either have no impact on the capsule structure or are not responsible for antigenic sites recognised by group 35 typing sera. Not surprisingly, most molecular tools using whole genome sequence data are not yet able to accurately serotype 33G (Supplementary Table S2). We have made the 33G *cps* sequences from this study publicly available (Genbank accession numbers OR509570-OR509589) so that *cps* databases can be updated. However, users of seroBA will not need to update the database as the 33G *cps* locus is already detected (called 10X). The US Centers for Disease Control and Prevention (CDC) method for qPCR serotyping (53) does not include primers for 10B or 33B and would therefore be unlikely to detect serotype 33G. DNA microarray designates 33G as 35A/10B-like due sequences in the 33G *cps* locus matching probes for some, but not all genes for the 35A and 10B *cps* loci. The TaqMan Array Card for pneumococcal serotyping (36) designates 33G as 10B due to the detection of *wcrD* in 10B, whereas the targets for other relevant serotypes such as 33B (*wciN*) and 35A (*wcrK*) are absent from the 33G *cps* locus. Serotypes 10B and 33B are not included in any licensed vaccines so it is unlikely 33G would be misidentified as a vaccine serotype in surveillance studies.

Serotype 33G has been found in healthy carriers ((54) and this study), in the nasopharynx of pneumonia patients (this study) as well as in invasive disease (16). These data indicate that 33G can not only colonize, but also cause disease. A total of 27 serotype 33G isolates have now been detected, including in Thailand (n=5) (18), South Africa (n=2) (16) and Mongolia (this study, n=20). We have tested over 18,000 swabs from across Asia and have not detected 33G in any of our previous pneumococcal carriage studies in healthy children or children with acute respiratory infection including Lao PDR (55), Fiji (56), Vietnam (57), Indonesia (58) or Papua New Guinea (59). It will be interesting to see whether 33G will be identified in other countries in the future, especially as sequence-based analysis becomes more accessible.

The discovery of serotype 33G has raised multiple questions that should form the basis of future research. Firstly, similar to the recently described serotype 33E (15), 33G lacks a branching monosaccharide, which are typically highly immunogenic (60). Thus, future research should focus on identifying whether production of a 33G capsule provides enhanced immune evasion capabilities or other biological advantages. Similarly, the function of *orf2* requires experimental investigation. Based on bioinformatic analysis, we propose *orf2* encodes a Wzy homolog replacing the ancestral *wzy* gene, which contains deletions and is likely non-functional. However, the role of ORF2 as the polymerase needs to be experimentally validated. Lastly, serotype 33G was only detected in Mongolia from 2018 onwards, following phased introduction of PCV13 from 2016. However, there are insufficient data to determine whether the emergence of 33G in Mongolia is related to vaccine introduction or not. Future studies should focus on gaining a better understanding of the geographic distribution of 33G, the prevalence of this serotype following vaccine introduction, as well as the potential of this serotype to cause pneumococcal disease particularly in the post-PCV era.

## MATERIALS AND METHODS

### Human ethics approval

Participants were recruited into either the carriage survey or pneumonia surveillance studies in accordance with the inclusion criteria described previously (19, 21, 22). Ethics approval was obtained from the Medical Ethics Review Committee at the Mongolian Ministry of Health and the Royal Children’s Hospital Human Research Ethics Committee (HREC reference numbers 33203 and 38045). Written consent was obtained by participants or their parents/caregivers before being included in the study.

### Pneumococcal identification and DNA microarray serotyping from nasopharyngeal swabs

Nasopharyngeal swabs were collected, stored, transported and tested following World Health Organization guidelines (37). Screening of swabs for pneumococci was conducted by *lytA* qPCR (24, 55). Culture of pneumococci from swabs was conducted on selective horse blood agar plates supplemented with 5 µg/ml gentamicin. DNA was then extracted from these cultures (QIAamp 96 DNA QIAcube HT kit) and tested by DNA microarray (Senti-SPv1.5 slides) as described previously (25) to determine serotype(s). Pneumococcal isolates of interest were purified and confirmed as pneumococci using standard identification tests (37) including optochin sensitivity and genotyping by MLST and GPSC.

### Whole genome sequencing

DNA was extracted from pneumococcal isolates using the QIAamp 96 DNA QIAcube HT kit as described previously (25). Libraries were constructed using the Illumina DNA Prep kit (Illumina). Sequencing was performed using NovaSeq as 2 x 150 bp paired end. Read quality was assessed with FastQC (https://www.bioinformatics.babraham.ac.uk/projects/fastqc/) with low quality reads removed using Trimmomatic (61). Genomes were assembled *de novo* using Spades 3.15.4 (62). Genome annotation was conducted with RAST (29). The serotype 33G *cps* sequences have been deposited in Genbank (accession numbers OR509570-OR509589)

### Molecular serotyping of 33X isolates

Quellung serotyping was performed as described previously (38). Genome assemblies or Fastq reads of 33X isolates were run through whole genome sequencing-based serotyping tools PneumoCaT (31), seroBA (32), SeroCall (33), PneumoKITy (34) and PfaSTer (35) using default parameters. DNA microarray was performed as described above. Pneumococcal serotyping using the TaqMan Array Card (SPN_CDC_V1 TAC) provided by Streptococcus Laboratory, Centers for Disease Control and Prevention and was performed as previously described (36).

### Competitive ELISA

Serum was generated by administering 2x10^3^ CFU of 33X pneumococci (strain PMP1486) to five-day old C57BL/6 mice intranasally without anaesthesia as described previously (63). At seven weeks post-infection, blood was collected by cardiac puncture, allowed to clot overnight at 4°C and subjected to centrifugation to collect serum. Animal work was approved by the Murdoch Children’s Research Institute (MCRI) Animal Ethics Committee (protocol A945) in accordance with the Australian code for the care and use of animals for scientific purposes (64). To prepare bacterial stocks for the ELISA, pneumococcal strains of serotypes 33X (PMP1486), 10B (PMP738), 33B (PMP790) and 14 (PMP829) were grown to OD 0.85-0.95 in THY broth (3% [w/v] Todd-Hewitt broth, 0.5% [w/v] yeast extract) at 37°C and 5% CO_2_ and stored in 80% (v/v) glycerol until required. Wells of high-binding Nunc Maxisorp plates were coated with PBS-washed 33X pneumococci (100 µl per well) and incubated overnight at 4°C. Competition ELISAs were conducted using a method adapted from previously established approaches (65, 66). Briefly, serum was incubated with PBS-washed preparations of pneumococci of different serotypes (33X, 10B, 33B or 14) in 10% (v/v) foetal calf serum (FCS) overnight at 4°C. Serum was diluted to 1:1000 in these reactions, which was determined to be the optimal dilution for high inhibition. The 96-well plates were blocked by adding 10% FCS for 1 h at 37°C. After washing with PBS containing 0.05% (v/v) tween, 50 µl of sera was added to the well and incubated for 1 h at 37°C and washed again. Biotinylated goat anti-mouse IgG (diluted 1:500 in 10% FCS) (Sigma-Aldrich) was added to the wells and incubated for 1 h at 37°C. Wells were washed, streptavidin-Horseradish Peroxidase conjugate (diluted 1:500 in 10% FCS) added and incubated for 1 h at 37°C. A 1:1 mixture of KPL TMB Peroxidase Substrate and KPL Peroxidase Substrate Solution B (Sera Care) was added to each well (50µl) and incubated in the dark for ∼12 min. The reaction was then stopped with the addition of 1 M H_3_PO_4_ and optical density read at 450 nm (and 630 nm as a reference) to calculate the percentage of inhibition.

### Extraction of 33X capsular polysaccharide

Pneumococcal strains were streaked from frozen stocks onto Columbia sheep blood agar plates and incubated overnight at 37°C and 5% CO_2_. Resultant colonies were then inoculated into modified Lacks medium (67) supplemented with 20 mM glucose and grown to an optical density at 600 nm of 0.5. Cultures were subjected to centrifugation at 3200 x g in a Heraeus centrifuge (Thermo Fisher Scientific, USA) and washed with ice-cold ultrapure water (in-house product). After another centrifugation, the supernatants were removed and pellets resuspended again in ultrapure water. Buffer-saturated phenol (Thermo Fisher Scientific, USA) with final concentration of 1% (v/v) was added to the suspension and incubated at room temperature overnight. Incubated samples were subjected to centrifugation for 30 min with 3200 x g at 4°C and supernatants collected. The collected supernatants were incubated with 3 µl benzonase nuclease (Sigma-Aldrich, USA) overnight at 37°C. Then 20 mg/ml Proteinase K solution (Roche, Switzerland) was added and again incubated overnight at 37°C. Next, the solution was transferred to a Millipore Amicon Ultra 30 kDa cut off membrane centrifugal filter unit (Merck Millipore, USA) (prewashed twice to remove glycerol) and subjected to centrifugation for 25 min at 3200 x g. The remaining solution above the filter was collected and evaporated under reduced pressure (25 mbar). Dried polysaccharide samples were submitted for chemical and NMR analysis.

### Sugar composition and linkage analysis

Composition analysis was carried out by chemical derivatisation of the sample both as alditol acetates and trimethylsilyl methyl glycosides. Alditol acetates were obtained after hydrolysis with 2 M trifluoroacetic acid (TFA) for 1 h at 125°C, followed by reduction with sodium borohydride (NaBD_4_) and peracetylation with acetic anhydride (68). Trimethylsilyl methyl glycosides were obtained by derivatization with the reagent Sylon™ HTP (Merck) after methanolysis of the polysaccharide with 3 M HCl in methanol at 85°C for 16 h (69). To determine the position of the glycosidic linkages, the polysaccharide 33X was permethylated following the protocol developed by Harris (70), hydrolyzed with 2 M TFA for 1 h at 125°C, reduced with sodium borodeuteride (NaBD_4_) and peracetylated with acetic anhydride to give a mixture of partially methylated alditol acetates.

The derivatised samples were analysed by GLC using an Agilent Technologies 6850 gas chromatograph equipped with a flame ionisation detector, using He as the carrier gas and a Zebron ZB-5 MSi capillary column (Phenomenex, 30 m x 250μm x 0.25μm). The following temperature programs were used: for alditol acetates, 3 min at 150°C, 150 - 270°C at 3°C/min, 2 min at 270°C; trimethylsilyl methyl glycosides, 1 min at 150°C, 150 - 280°C at 5 °C/min, 2 min at 280°C; for partially methylated alditol acetates, 1 min at 90°C, 90 - 140°C at 25°C/min, 140 - 200°C at 5°C/min, 200 - 280 °C at 10°C/min, 10 min at 280°C. GLC-MS analyses were carried out on an Agilent Technologies 7890 A gas chromatograph coupled to an Agilent Technologies 5975C VL MSD using the same column and the temperature programs of the GLC analyses. Values of the integrated area of the partially methylated alditol acetates were corrected by the effective carbon response factors (71).

### NMR analysis

A sample of polysaccharide 33X (1-2 mg) was dissolved in deuterium oxide (D_2_O) (Sigma-Aldrich, USA), subjected to centrifugation for 5 min at 16800 x g, transferred to 1.7 mm NMR tubes (Bruker, USA) and submitted for NMR analysis. Initial studies were conducted on a Bruker Neo 500 MHz NMR spectrometer equipped with a 1.7 mm TXI probehead; the probe temperature was set at 25°C. Further NMR experiments were performed after recovery of the sample, cycles of D_2_O exchange and transfer to a 5 mm tube, on a Bruker Avance III 600 MHz NMR spectrometer equipped with a BBO Prodigy cryoprobe and processed using standard Bruker software (Topspin 3.2). The probe temperature was set at 60°C or 70°C. 1D (^1^H, ^31^P and ^13^C) and 2D, COSY, TOCSY, NOESY, HSQC and HMBC NMR experiments were performed. 2D COSY and NOESY experiments were recorded with pre-saturation of HOD, whereas the TOCSY experiments were performed using DOSY to remove signals from low molecular weight components (ledbpgpml2s2d). 2D TOCSY experiments were recorded using a mixing time of 180 ms and the 1D variants using 180 or 200 ms. 2D NOESY experiments were recorded using a mixing time of 300 ms and the 1D variants using 300 or 500 ms. The HSQC experiment was optimized for J = 145 Hz (for directly attached ^1^H-^13^C correlations), and the HMBC experiments optimized for a coupling constant of 6 Hz (for long-range ^1^H-^13^C correlations) and 10 Hz (for ^1^H-^31^P correlations). To improve sensitivity by performing many scans, the 2D experiments were recorded using non-uniform sampling: 40% for homonuclear and 20-30% for heteronuclear experiments. Spectra were referenced to residual CWPS: ^1^H signal at 3.23 ppm, ^13^C signal at 54.5 ppm and the shielded ^31^P signal at 1.30 ppm (72).

## ACKNOWLEDGEMENTS

We thank all the staff involved in the recruitment and nasopharyngeal swab collection in the carriage surveys and pneumonia surveillance programs in Mongolia as well as the Translational Microbiology Group at the Murdoch Children’s Research Institute. We also thank all the participants and their families involved in these studies. Thank you to Beth Temple for epidemiological advice regarding nasopharyngeal swab data. We thank Lesley McGee and Srinivasan Velusamy for providing the pneumococcal TaqMan Array Cards.

## AUTHOR CONTRIBUTIONS

Conceptualization: SM, CS

Methodology and study design: SM, JH, FMR, EKM, OT, TM, CVM, EMD, BDG, PVL, PC, NR, MH, CS

Data analysis: SM, CLP, PVL, PC, NR, MH, CS

Experimental investigation: SM, BDO, BB, IG, JPW, CLP, EN, SWL, JH, SDB, NR

Funding acquisition: SM, EKM, FMR, MH, CS

Writing of the original manuscript draft: SM, MH, NR, CS

Review and editing of the manuscript for submission: SM, BDO, JPW, BB, CLP, EN, IG, SWL, JH, SDB, FMR, EKM, OT, EMD, BDG, TM, CVM, PVL, PC, NR, MH, CS

