## Supplementary data for "*Streptococcus pneumoniae* serotype 33G: genetic, serological, and structural analysis of a new capsule type"

```
Y5914-11_acyltransferase      ATGAAGAAGGAATATGATATTTTAAAAGTAATTGCCATTTTAATGGTTGTATTAAGCCAC	60
33X_orf3/4region              ATGAAGAAGGAATATGATATTTTAAAAGTAATTGCCATTTTAATGGTTGTATTAAGCCAC	60
                              ************************************************************

Y5914-11_acyltransferase      AGTACATACTATGTGATTTCGACTAAGTACGGGGGGGTTGATTATCAACAATATATAAAT	120
33X_orf3/4region              AGTACATACTATGTGATTTCGACTAAGTACGGGGGGA----TTATCAACAATATATAAAT	116
                              ************************************.    *******************

Y5914-11_acyltransferase      CAAAATTTATCGTTGGTATTGTATAAAGTTTTTGATAAAGTAAGAGAAGTATTATATTAC	180
33X_orf3/4region              CAAAATTTATCGTTGGTATTGTATAAAGTTTTTGATAAAGTAAGAGAAGTATTATATTAC	176
                              ************************************************************

Y5914-11_acyltransferase      TTCCATATGCCACTTTTTATGGCATTGTCAGGAGCTTTCTACTATCTTCAGGTTCAAAGA	240
33X_orf3/4region              TTCCATATGCCACTTTTTATGGCATTATCAGGAGCTTTCTACTATCTTCAGGTTCAAAGA	236
                              **************************.*********************************

Y5914-11_acyltransferase      GATAAATGGTCTACTTTGAAATTAATTGTGCAAAA-----TAAGGCAAAGAGATTGCTTT	295
33X_orf3/4region              GATAAATGGTTTACTTTAAAATTAATTGTGCAAAATAAAATAAGGTAAAGAGATTGCTTT	296
                              ********** ******.*****************     ***** **************

Y5914-11_acyltransferase      TTCCTTTCATTATATTTACTATTCTTTATTCAATACCAATAAAATATATTTCAAATTATT	355
33X_orf3/4region              TTCCTTTCATTATATTTACTGTCCTTTATTCAATACCAATAAAATATATTTCAAATTATT	356
                              ********************.* *************************************

Y5914-11_acyltransferase      TTGATTCTACAGATCCTTTTAAAGCATTTGTAGGACAATTTTTCTTAATTGGGAATAGTC	415
33X_orf3/4region              TTGATTTTACAGCTCCTTTTAAAGCATTTGTAGGAGAATTTTTCTTAATTGGAAATAGTC	416
                              ****** *****.********************** ****************.*******

Y5914-11_acyltransferase      ATTTATGGTATTTATATGCTTTATTTATTATTTTTATATTTGCATTCTATACGCTAAAAA	475
33X_orf3/4region              ATTTATGGTATTTATATGCTTTATTTATTATTTTTATATTTGCATTCTATACGCTAAA-A	475
                              ********************************************************** *

Y5914-11_acyltransferase      AAGAAACAAAGCTTGCTACTTATGTTGTCTTTTATGTTCTGCATATTTTGAGTTACAAAA	535
33X_orf3/4region              AAGAAACAAATCTTGCTACTTATGTTGTTTTTTATGTTCTGTATATTTTGAGTTACAAAA	535
                              ********** ***************** ************ ******************

Y5914-11_acyltransferase      TAGAACTCCCACTATTTAAAGTACCTCTTCAATTTTTATTTTACTTTAGTTTAGGCTTTT	595
33X_orf3/4region              TAGAACTCACACTATTTAAAGTACCTCTTCAATTTTTATTTTACTTTAGTTTAGGCTTCT	595
                              ********.************************************************* *

Y5914-11_acyltransferase      TGTTTGAATCAAATAGAGAAAAATATAATCAATTTATTAATAAGAAAAAAAATTATGTTT	655
33X_orf3/4region              TGTTTGAATCAAATAGAGAAAAATATAATCAATTTATTAATAGGAAA-AAAATTATATTT	654
                              ******************************************.**** ********.***

Y5914-11_acyltransferase      TGTTATTATCTACGGTATTTGCCTTAATGGTTTTATTAAATCTCTATGTTAAGTTTACTC	715
33X_orf3/4region              TGTTATTATCTACGGTATTTGTCTTAATGGTTTTG-------------TTAAATTTACTC	701
                              ********************* ************.             ****.*******

Y5914-11_acyltransferase      AGGTATTGTTTAGTAAAGTATTAGTTGAGTTAATGGCTGTCTTGGGTTCATTATTAACGT	775
33X_orf3/4region              AGGTATTTTTTAGTAAAGTATTAGTTGAGCTATTGGCTGTCTTGGGTTCATTATTAACGT	761
                              ******* ********************* **:***************************

Y5914-11_acyltransferase      ATAGCATCGCGTATCAATTATCTCAAAAGAAAAGTGTTGGTGATGCTAGTATTTTTAAGA	835
33X_orf3/4region              ATAGCATCGCATATCAATTATCTCAAAAGAAAAGTGTTGGTGATGCTAGTATTTTTAAGA	821
                              **********.*************************************************

Y5914-11_acyltransferase      TGATTCTAATTAATGGATTAGGTATATATATTTTTTCTGATCCATTAAATTATTTAATCT	895
33X_orf3/4region              TGATTCTAATTAATGGATTAGGTATATATATTTTTTCTGATCCATTAAATTATTTAATCT	881
                              ************************************************************

Y5914-11_acyltransferase      TGAAATTAAGTTATTCCCTAAATTCGTATTTTATGTTTACACCATTAGGAATAGTTGTTT	955
33X_orf3/4region              TGAAGTTAAGTTATTCCCTAAATTCGTATTTTATGTTTACACCAGTAGGAATAATTGTTT	941
                              ****.*************************************** ********.******

Y5914-11_acyltransferase      TAGTGGTGCTACGATTTTTTCTTACGTTATTTATATCATTGATAGGAACAATTATATTTA	1015
33X_orf3/4region              TAGTAGTATTACGATTTTTTCTCACGTTATTCATATCATTGATAGGAACAATCATATTT-	1000
                              ****.**. ************* ******** ******************** ******

Y5914-11_acyltransferase      AAAAATATTTTAAAAAATACAAATGGCTAGTCAACTAG----------------------	1053
33X_orf3/4region              ----------AAAAAAATACAAATGGCTAGTCAACTAGGTAGAGACAATAGCTATTTAAA	1050
                                        :***************************

Y5914-11_acyltransferase      -------	1053
33X_orf3/4region              AAGTTAA	1057
```

Figure S1: DNA sequence alignment of the acyltransferase gene from *S. oralis* subsp. *Dentisani* strain Y5914-11 (Genbank accession NZ_NCUW01000028, locus tag B7709_03985) with the *orf3-orf4* region from the pneumococcal 33X *cps* locus. Red and blue text denote the predicted *orf3* and *orf4* sequences detected in the 33X *cps* locus. Nucleotides highlighted in yellow denote stop codons created as a result of frameshift mutations.

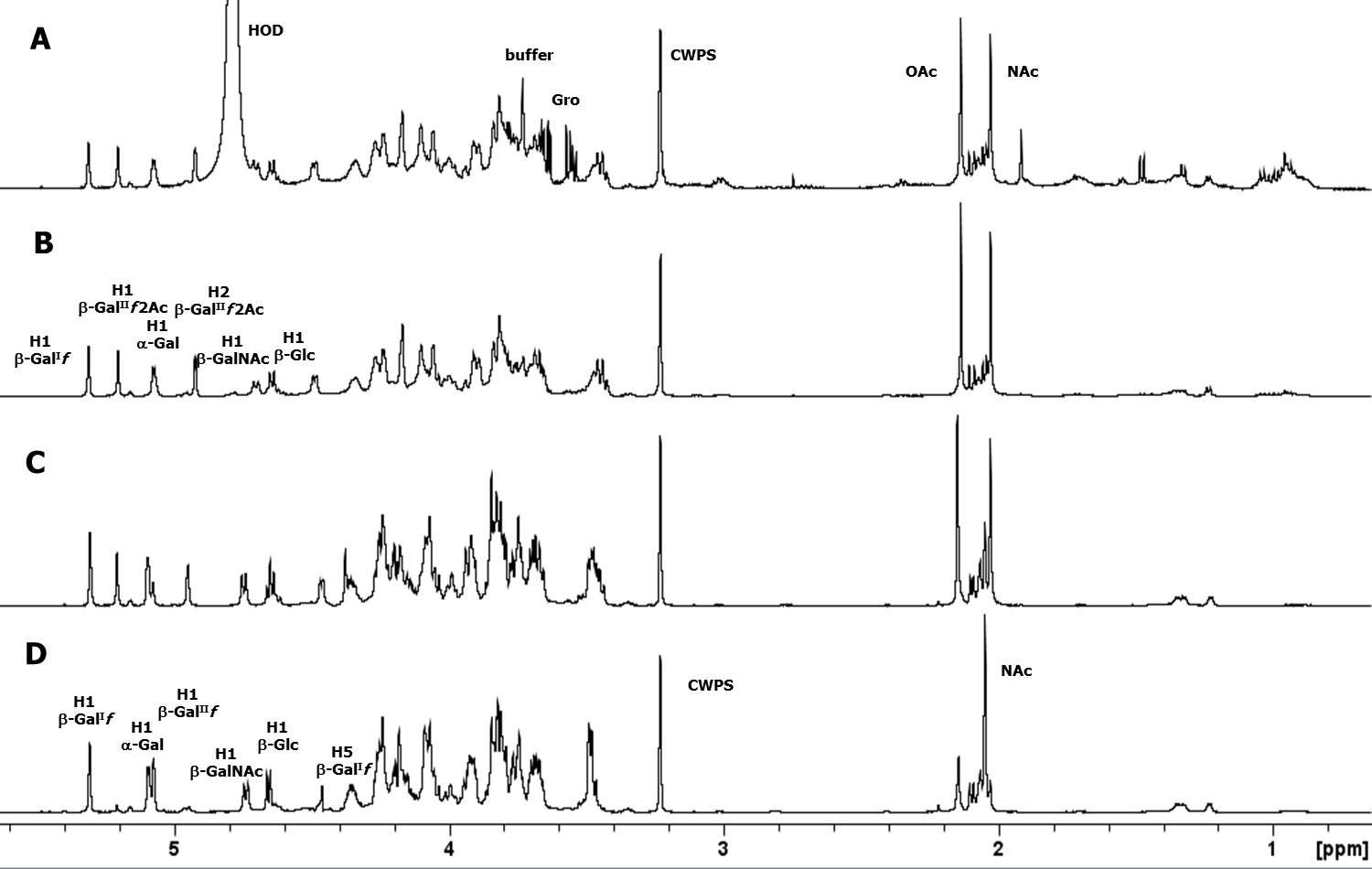

Figure S2: 1D proton NMR spectra of pneumococcal serotype 33G polysaccharide, some diagnostic anomeric and ring protons are labeled. (A) 1D 1H NMR at 500 MHz and 25°C, (B) 1D DOSY at 500 MHz and 25°C, (C) 1D DOSY at 600 MHz and 70°C (partially de-O-acetylated), and (D) 1D DOSY at 600 MHz and 60°C (de-O-acetylated polysaccharide). The 1D DOSY (Figure 3D) gave five anomeric signals of similar intensity at *δ* 5.31 (br s), 5.09 (d, ^3^*J*_H-1,H-2_ = 3.6 Hz), 5.07 (br s), 4.74 (d, ^3^*J*_H-1,H-2_ = 8.4 Hz) and 4.66 (d, ^3^*J*_H-1,H-2_ = 7.8 Hz) consistent with a hexasaccharide repeat unit. The broad singlets were attributed to β-galactofuranose residues (confirmed by the presence of characteristic deshielded C1 resonances at 108.5 and 109.3 ppm, Figure S4), whereas the chemical shift and vicinal coupling constant values of the remaining signals were in a good agreement with an α-pyranose and two β-pyranose sugars.

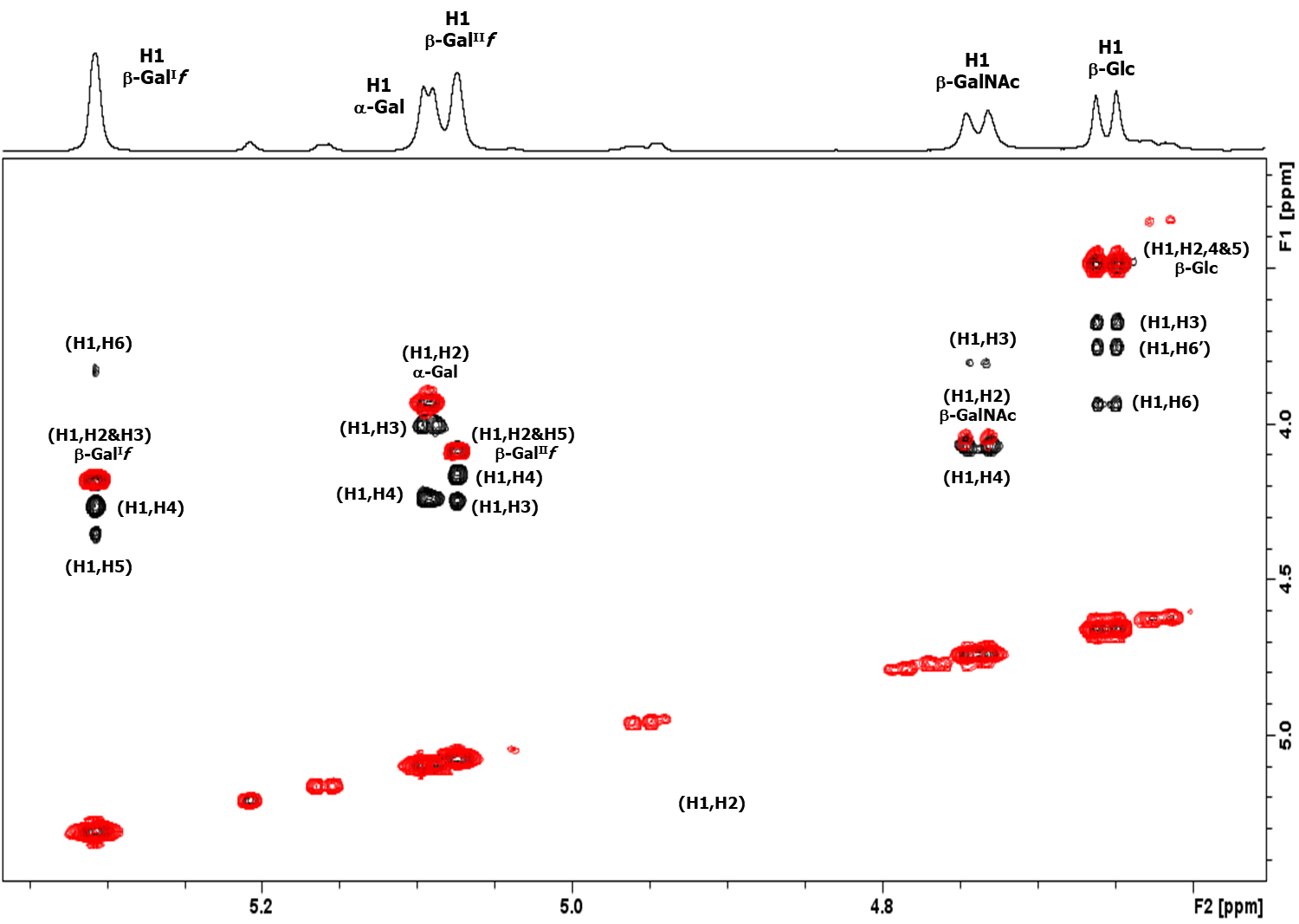

Figure S3: The 2D ^1^H-^1^H overlay for polysaccharide 33G: COSY (red)/ TOCSY (black) recorded at 600 MHz (60°C). The major crosspeaks from H1 for the 33G repeat unit are labeled. The DOSY-TOCSY experiment was recorded using a mixing time of 180 ms. The COSY spectrum gave H1 to H2 for each residue and the TOCSY experiment established further correlations for each spin system depending on the sugar type: H1 to H6 for β-Gal^I^*f*, H1 to H4 for α-Gal*p*, H1 to H5 for β-Gal^II^*f*, H1 to H4 for β-GalNAc and H1 to H6 for β-Glc. The assignments were confirmed, and some additional protons identified using 1D TOCSY profiles (Figure S3).

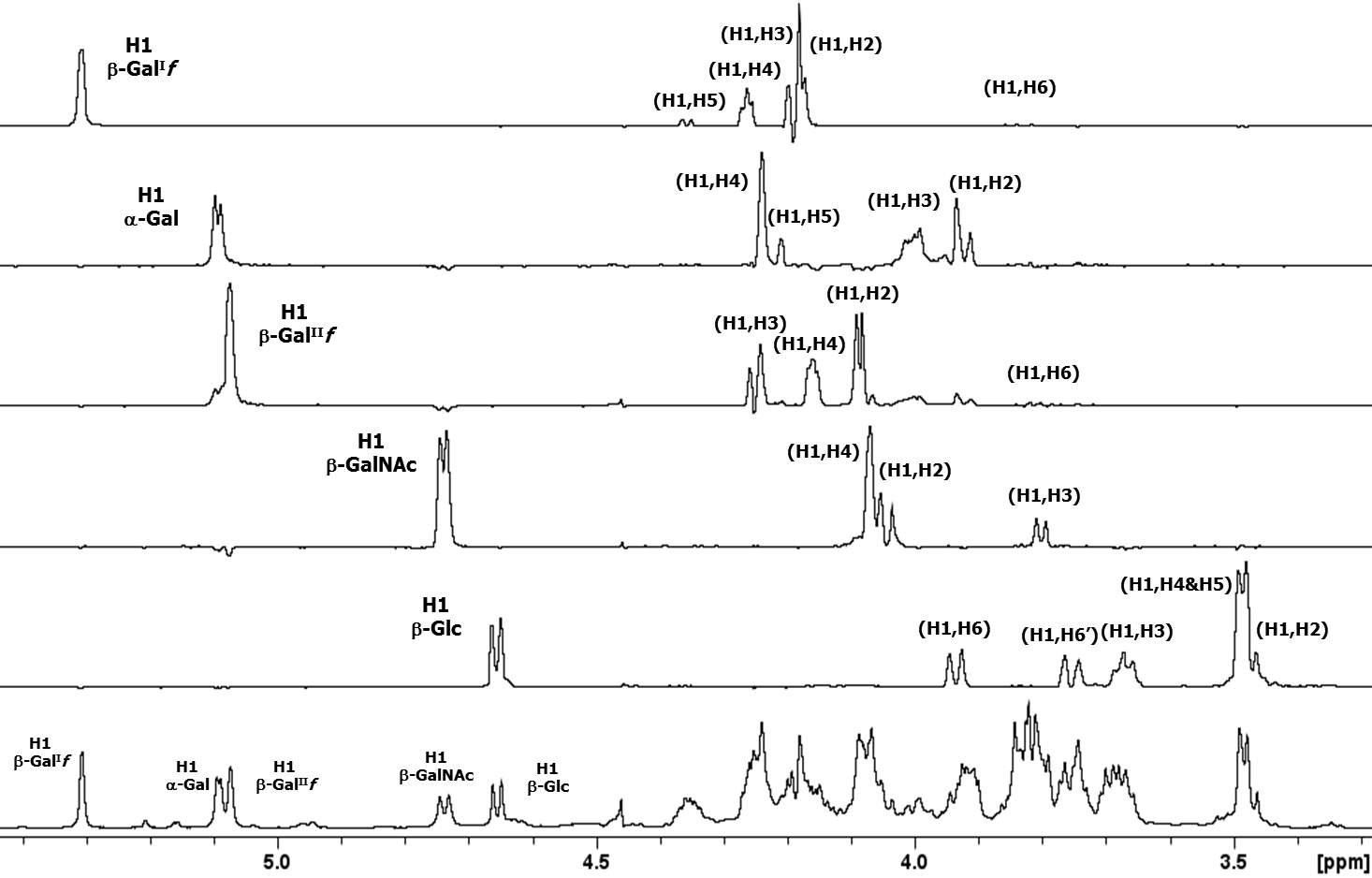

Figure S4: Overlay of 1D TOCSY (180 ms) profiles extracted from the 2D DOSY-TOCSY experiment overlaid over the 1D DOSY spectrum. The spectra corroborated the 2D NMR assignments (Figure S2).

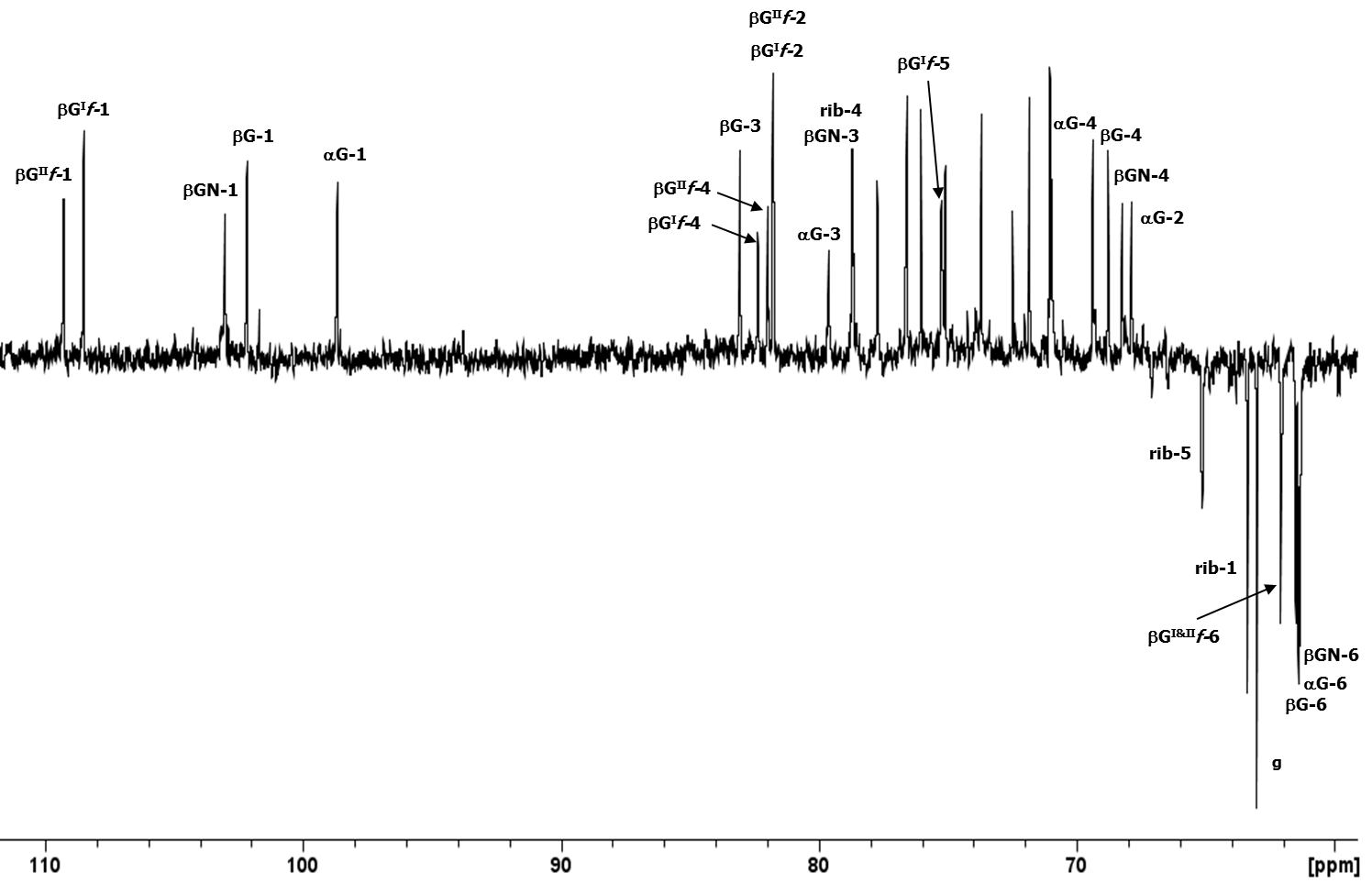

Figure S5: Expansion of the ^13^C DEPT-135 spectrum of polysaccharide 33G recorded at 150 MHz and 60°C. Diagnostic peaks from the hexasaccharide repeating unit have been labeled (βG^I^*f* = β-Gal^I^*f*, αG = α-Gal*p*, βG^II^*f* = β-Gal^II^*f*, βGN = β-GalNAc, β-G = β-Glc and rib = ribitol). Additional peaks are due to glycerol (g). The presence of ribitol-5- phosphate was confirmed by the methylene peaks at *δ* 63.4 (H1s at 3.83 and 3.68 ppm) and 65.1 ppm (H5s at 4.21 and 4.14 ppm) and splitting of the latter by coupling with ^31^P (*J*_H,P_ = 5.5 Hz).

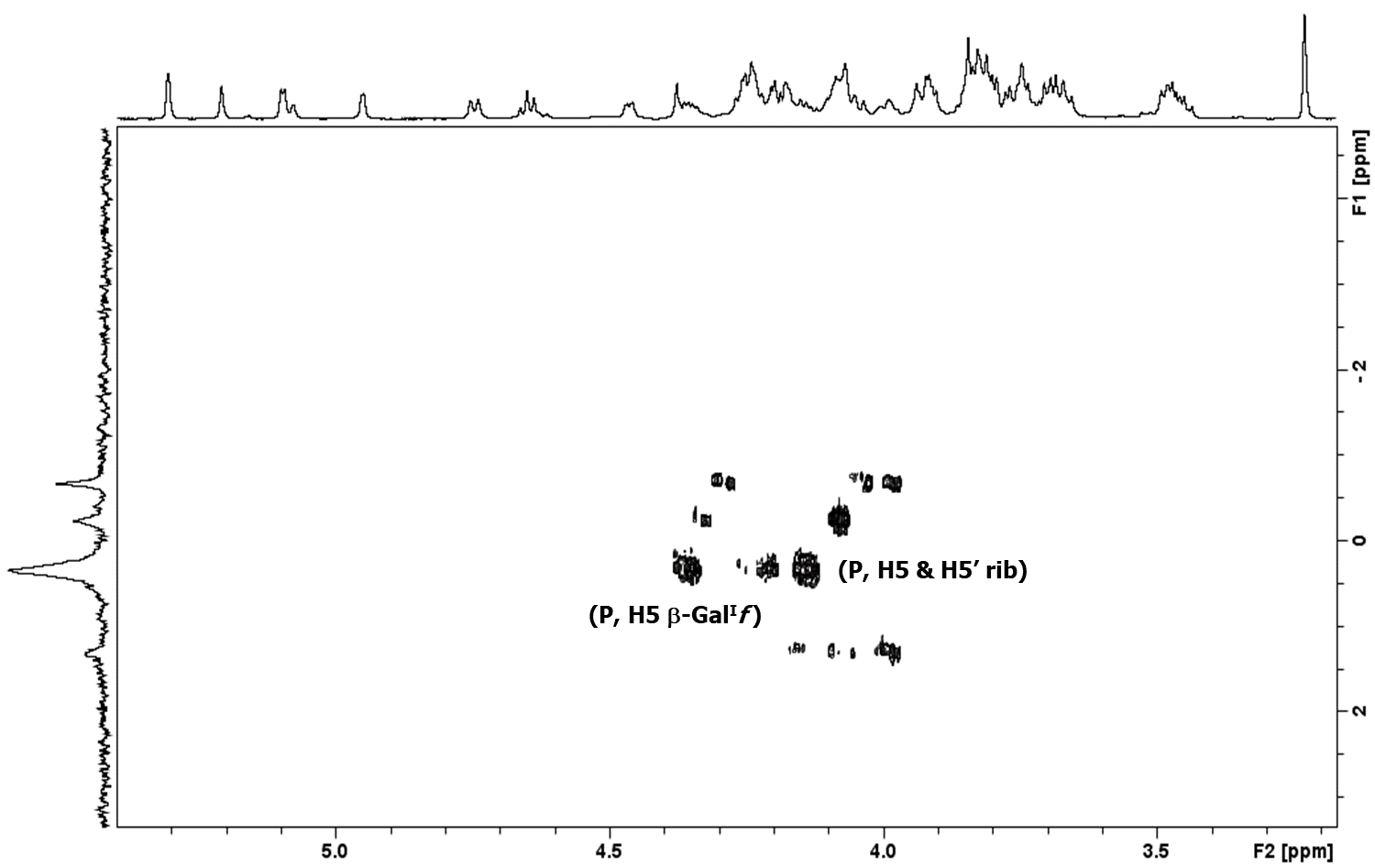

Figure S6. The 2D ^1^H–^31^P HMBC spectrum for polysaccharide 33G recorded at 600 MHz (70°C). The major crosspeaks from the phosphodiester at 0.34 ppm are labeled (H5s of ribitol and H5 of β-Gal^I^*f*); the additional unlabeled peaks are from residual CWPS. The HMBC experiment was optimized for a coupling constant of 10 Hz.

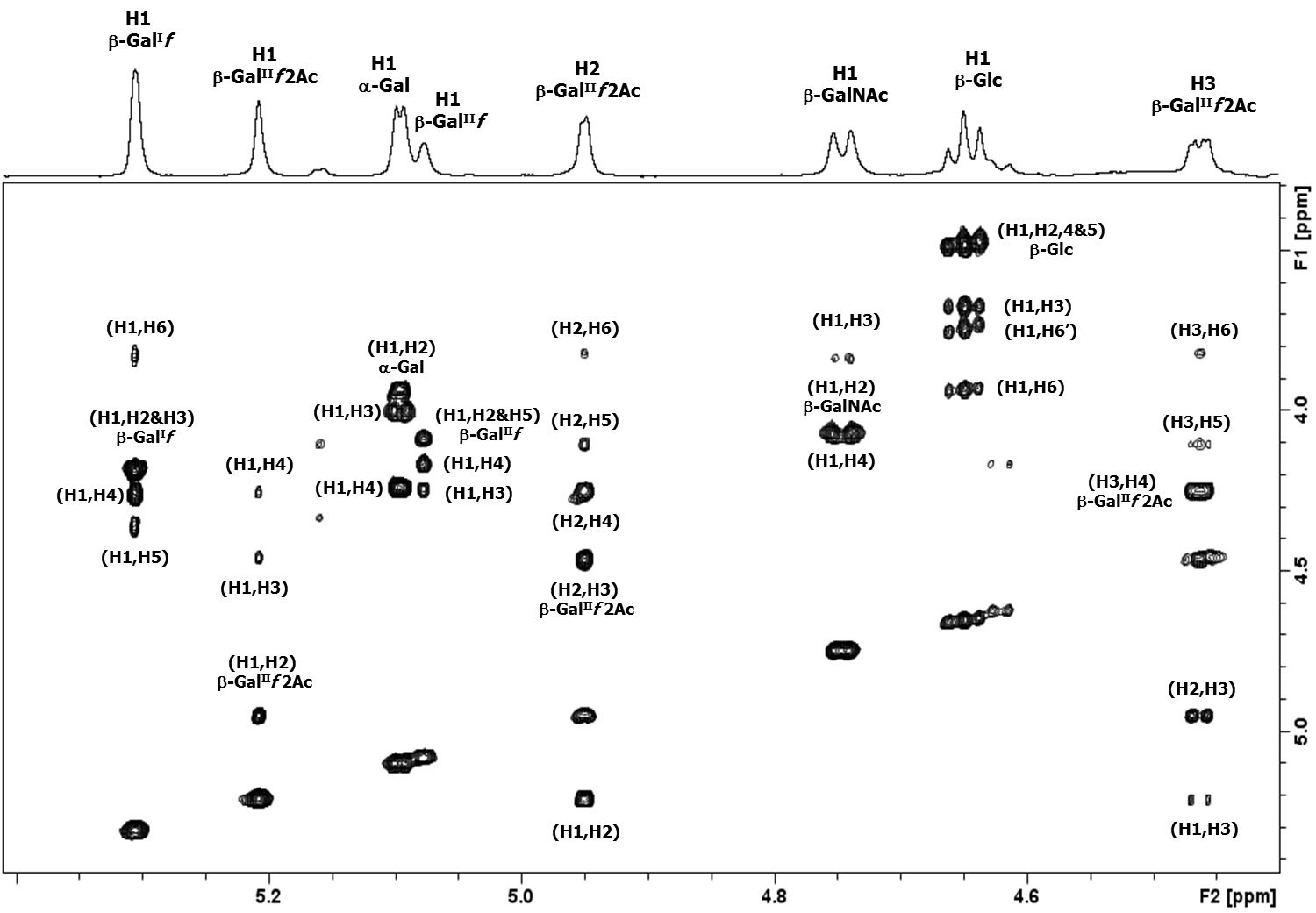

Figure S7: The 2D ^1^H-^1^H TOCSY of the partially O-acetylated 33G polysaccharide recorded at 600 MHz (70°C). The major crosspeaks from H-1 and other diagnostic peaks in the anomeric region for the 33G repeat unit are labeled.

Table S1. Pneumococcal-positive nasopharyngeal swabs containing 33X pneumococci in Mongolia

| **Year** | **Pneumococcal-positive swabs containing serotype 33X**  **(n/N, %)** |
| --- | --- |
| 2015 | 0/701 (0%) |
| 2016 | 0/600 (0%) |
| 2017 | 0/1006 (0%) |
| 2018 | 3/688 (0.4%) |
| 2019 | 6/982 (0.6%) |
| 2020 | 7/462 (1.5%) |
| 2021^a^ | 0/33 (0%) |
| 2022 | 4/328 (1.2%) |

^a^ Overall sample numbers in 2021 were very low consistent with the non-pharmaceutical interventions in place for the COVID-19 pandemic

Table S2. Summary of molecular serotyping results for 33X pneumococci

|  | **DNA microarray** (1) | **PneumoCaT** (2) | **seroBA** (3) | **serocall** (4) | **PneumoKITy** (5) | **PfaSTer** (6) | **TaqMan Array Card** (7) |
| --- | --- | --- | --- | --- | --- | --- | --- |
| **Serotype result** | 35A/10B-like^a^ | Non-typeable | 10X | Non-typeable | Non-typeable | Non-typeable (Mean probability = 0.17) | 10B |
| **Best serotype match** | N/A | 35B  (Mean coverage = 59%) | N/A | N/A | 35B  (Mean hash score = 44%) | 35D  (low confidence) | N/A |

^a^a ‘like’ designation indicates that DNA microarray could not designate to a known serotype, but has sequence similarity to portions of the serotype 35A and 10B *cps* loci
